# DeepSide: A Deep Learning Framework for Drug Side Effect Prediction

**DOI:** 10.1101/843029

**Authors:** Onur Can Uner, Ramazan Gokberk Cinbis, Oznur Tastan, A. Ercument Cicek

**Affiliations:** Department of Computer Engineering, Bilkent University, Ankara, Turkey 06800; Department of Computer Engineering, Middle East Technical University, Ankara, Turkey 06800; Faculty of Engineering and Natural Sciences, Sabanci University, Istanbul, Turkey 34956; Computational Biology Department, Carnegie Mellon University, Pittsburgh, PA 15213

## Abstract

Drug failures due to unforeseen adverse effects at clinical trials pose health risks for the participants and lead to substantial financial losses. Side effect prediction algorithms have the potential to guide the drug design process. LINCS L1000 dataset provides a vast resource of cell line gene expression data perturbed by different drugs and creates a knowledge base for context specific features. The state-of-the-art approach that aims at using context specific information relies on only the high-quality experiments in LINCS L1000 and discards a large portion of the experiments. In this study, our goal is to boost the prediction performance by utilizing this data to its full extent. We experiment with 5 deep learning architectures. We find that a multi-modal architecture produces the best predictive performance among multi-layer perceptron-based architectures when drug chemical structure (CS), and the full set of drug perturbed gene expression profiles (GEX) are used as modalities. Overall, we observe that the CS is more informative than the GEX. A convolutional neural network-based model that uses only SMILES string representation of the drugs achieves the best results and provides 13.0% macro-AUC and 3.1% micro-AUC improvements over the state-of-the-art. We also show that the model is able to predict side effect-drug pairs that are reported in the literature but was missing in the ground truth side effect dataset. DeepSide is available at http://github.com/OnurUner/DeepSide.

## 1 Introduction

Computational methods hold great promise for mitigating the health and financial risks of drug development by predicting possible side effects before entering into the clinical trials. Several learning based methods have been proposed for predicting the side effects of drugs based on various features such as: chemical structures of drugs [25, 1, 23, 8, 19, 34, 17, 9, 2, 5], drug-protein interactions [35, 33, 8, 19, 34, 17, 37, 2, 15, 36], protein-protein interactions (PPI) [8, 9], activity in metabolic networks [38, 26], pathways, phenotype information and gene annotations [8]. In parallel to the above mentioned approaches, recently, deep learning models have been employed to predict side effects: (i) [31] uses biological, chemical and semantic information on drugs in addition to clinical notes and case reports and (ii) [4] uses various chemical fingerprints extracted using deep architectures to compare the side effect prediction performance.

While these methods have proven useful for predicting adverse drug reactions (ADRs – used interchangeably with drug side effects), the features they use are solely based on external knowledge about the drugs (i.e., drug-protein interactions, etc.) and are not cell or condition (i.e., dosage) specific. To address this issue, Wang et al. (2016) utilize the data from the LINCS L1000 project [32]. This project profiles gene expression changes in numerous human cell lines after treating them with a large number of drugs and small-molecule compounds. By using the gene expression profiles of the treated cells, [32] provides the first comprehensive, unbiased, and cost-effective prediction of ADRs. The paper formulates the problem as a multi-label classification task. Their results suggest that the gene expression profiles provide context-dependent information for the side-effect prediction task. While the LINCS dataset contains a total of 473,647 experiments for 20,338 compounds, their method utilizes only the highest quality experiment for each drug to minimize noise. This means that most of the expression data are left unused, suggesting a potential room for improvement in the prediction performance. Moreover, their framework performs feature engineering by transforming gene expression features to enrichment vectors of biological terms. In this work, we investigate whether the incorporation of gene expression data along with the drug structure data can be leveraged better in a deep learning framework without the need for feature engineering.

In this study, we propose a deep learning framework, DeepSide, for ADR prediction. DeepSide uses only (i) *in vitro* gene expression profiling experiments (GEX) and their experimental meta data (i.e., cell line and dosage - META), and (ii) the chemical structure of the compounds (CS). Our models train on the full LINCS L1000 dataset and use the SIDER dataset as the ground truth for drug - ADR pair labels [13]. We experiment with five architectures: (i) a multi-layer perceptron (MLP), (ii) MLP with residual connections (ResMLP), (iii) multi-modal neural networks (MMNN.Concat and MMNN.Sum), (iv) multi-task neural network (MTNN), and finally, (v) SMILES convolutional neural network (SMILESConv).

We present an extensive evaluation of the above-mentioned architectures and investigate the contribution of different features. Our experiments show that CS is a robust predictor of side effects. The base MLP model, which uses CS features as input, produces ~11% macro-AUC and ~2% micro-AUC improvement over the state-of-the-art results provided in [32], which uses both GEX (high quality) and CS features. The multi-modal neural network model, which uses CS, GEX and META features and uses summation in the fusion layer (MMNN.Sum) achieves 0.79 macro-AUC and 0.877 micro-AUC which is the best result among MLP based approaches. We also find out that when the chemical structure features are fully utilized in a complex model like ours, it overpowers the information that is obtained from the GEX dataset. The convolutional neural network that only uses the SMILES string representation of the drug structures achieves the best result among all the proposed architectures with provides 13.0% macro-AUC and 3.1% micro-AUC improvement over the state-of-the-art algorithm. Finally, inspecting the confident false positives predictions reveal side effects that are not reported in the ground truth dataset, but are indeed reported in the literature. DeepSide is implemented and released at http://github.com/OnurUner/DeepSide.

## 2 Methodology

### 2.1 Problem Formulation

The problem of side effect prediction is modelled as a multi-label classification task. For a given drug *i*, the target label is a binary vector, **y**_*i*_ = [*y*_*i*,1_, *y*_*i*,2_, …, *y*_*i,d*_], where *d* is the number of side effects and *y*_*i,j*_ = 1 indicates that the drug *i* has side effect *j*; *y*_*i,j*_ = 0 indicates otherwise. Our dataset contains *n* samples (drugs), each represented by a pair of drug feature vector **x**_**i**_ and an accompanying side effect vector (classes) 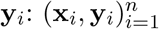.

### 2.2 Datasets

The LINCS L1000 dataset (GSE92742) contains the GEX profiles of 76 cell lines, treated with 20,413 small-molecule compounds [28]. There are 473,647 signature experiments that differ by the dosage, timing, and cell line (Level 5 data). In each experiment, the expression levels of 978 landmark genes are recorded. The study has two development phases: Phase 1 and Phase 2. Phase 1 contains approved drugs, whereas Phase 2 contains drugs that are at an experimental stage. To be able to compare our results with those in [32], we use Phase 1 data and process the dataset in the same manner. The authors report that their best result is obtained with the feature set that is a combination of gene ontology (GO) transformed gene expression profiles and chemical structures (CS). Their set of drugs with this feature set (GO + CS) contains 791 compounds. We use these 791 drugs to build our models. In total, there are 18,832 experiments for these 791 drugs in the LINCS L1000 dataset.

The META information for each of the 18,832 experiments from the LINCS project is also used as features. META information contains (i) the cell line on which the experiment is conducted on, (ii) the timing of the experiment, and (iii) dosage information. The meta information exists for 70 cell lines, 20 dosage levels and 3 time points (i.e. 6h, 24h, 48h). Note that for a given drug, the experiments do not cover all possible combinations of these conditions. META data is represented as one-hot encoding vectors. The corresponding feature vector has a length of 93. The total length of the concatenated GEX and META feature vectors is 1071. For all models, whenever META data is used, it is concatenated with the 978 landmark GEX features.

We obtain the drug side effect information (labels) from the SIDER Database [13] (downloaded on Feb 5, 2018). The side effects that are observed with fewer than ten drugs are excluded as also done in [32]. This filtering stage leaves us with 1052 side effects in total. In order to group side effects, we utilize the ADR ontology database (ADReCS), which provides a hierarchical classification of side effects in a four-level tree [3].

The CS features are encoded with OpenBabel Chemistry Toolbox [20] to create a 166-bit MACCS chemical fingerprint matrix for each drug (a binary vector of length 166). A SMILES string is an alternative representation for the 2D molecular graph of a drug/small molecule as a 1D string. The SMILES strings are downloaded from PubChem [11]. These are used to create the chemical fingerprints of the drugs for the 1D convolution used in SMILESConv model. RDKit Cheminformatics toolbox is used to extract extended SMILES Strings of the drugs [14]. The extended SMILES strings contain all the primary chemical bonds as well as the hydrogen bonding information explicitly. Zero-padding is used to have a uniform representation among all drugs. The alphabet contains 33 unique characters, including the end of sequence character. We further generate a pruned drug dataset to compare SMILESConv model with others. We filter out drugs with SMILES representation that have less than 100 characters and more than 400 characters. 615 out of 791 drugs pass this filtering step. For these drugs, we apply the additional filtering for removing side effects with less than ten drugs. In the end, 615 drugs and 1042 side effects pairs remain in this pruned dataset. Finally, we remove the characters that occur only once in all SMILES strings from the character vocabulary and replace them with underscore symbol.

### 2.3 The DeepSide Architectures

We propose the following deep learning architectures for ADR prediction: (i) a simple multi-layer perceptron, (ii) its residual variant, (iii) multi-modal network architectures that pre-transform inputs from each domain separately, (iv) multi-task neural network, and finally, (v) a convolutional neural network based approach for incorporating SMILES representation.

#### Multi-layer perceptron (MLP)

Our MLP [22] model takes the concatenation of all input vectors and applies a series of fully-connected (FC) layers. Each FC layer is followed by a batch normalization layer [10]. We use ReLU activation [16], and dropout regularization [27] with a drop probability of 0.2. The sigmoid activation function is applied to the final layer outputs, which yields the ADR prediction probabilities. The loss function is defined as the sum of negative log-probabilities over ADR classes, i.e. the multi-label binary cross-entropy loss (BCE). An illustration of the architecture for CS and GEX features is given in Figure 1.

**Fig. 1:**
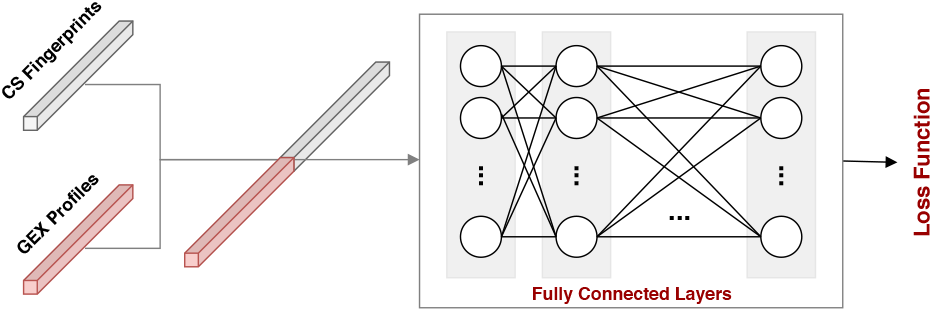
Our multi-Layer perceptron (MLP) architecture, which takes the concatenation of GEX and CS features.

#### Residual multi-layer perceptron (ResMLP)

The residual multi-layer perceptron (ResMLP) architecture is very similar to MLP, except that it uses residual-connections across the fully-connected layers. More specifically, the input of each intermediate layer is element-wise added to its output, before getting processed by the next layer. Such residual connections have been shown to reduce the vanishing gradient problem to a large extend [7]. This effectively allows deeper architectures, therefore, potentially learning more complex and parameter-efficient feature extractors.

#### Multi-modal neural networks (MMNN)

The multi-modal neural network approach contains distinct MLP sub-networks where each one extract features from one data modality only. The outputs of these sub-networks are then *fused* and fed to the classification block. For feature fusion, we consider two strategies: concatenation and summation. While the former one concatenates the domain-specific feature vectors to a larger one, the latter one performs element-wise summation. By definition, for summation based fusion, the domain-specific feature extraction sub-networks have to be designed to produce vectors of equivalent sizes. We refer to the concatenation and summation based MMNN networks as MMNN.Concat and MMNN.Sum, respectively. The MMNN.Concat approach is illustrated in Figure 2a.

**Fig. 2:**
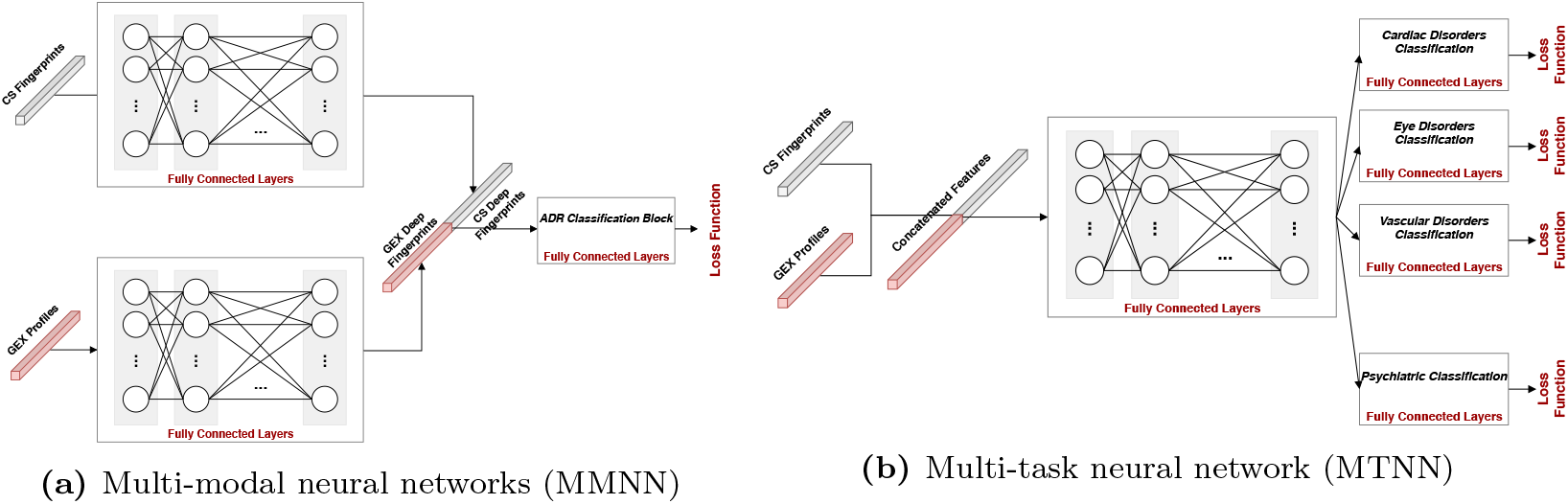
Multi-modal and Multi-task Neural Network architectures. (a) The concatenation variant of the multi-modal neural network (MMNN.Concat) architecture, which has two input branches for the GEX and CS features. The outputs of these networks are concatenated and fed into a fully connected multi-layer classification block. b) The multi-task neural network (MTNN) architecture, which learns a shared embedding for all class groups in the shared layers. The embedding is then fed into separate fully-connected multi-layer classification blocks for each class group, which learn task specific models.

#### Multi-task neural network (MTNN)

Our multitask learning (MTL) based architecture aims to take the side effect groups obtained from the taxonomy of ADReCS into account. For this purpose, the approach defines *shared* and *task-specific* MLP sub-network blocks. The shared block takes the concatenation of GEX and CS features as input and outputs a joint embedding. Each task-specific sub-network then converts the joint embedding into a vector of binary prediction scores for a set of inter-related side-effect classes.

We define 24 side-effect groups according to the ADR ontology (see Section 2.2). Here, a side effect is allowed to be a member of multiple groups. For instance, in the ADR ontology, *nausea* is grouped under both *stomach disorders* and *dizziness* sub-groups. For such side effects, our model will output more than one probability estimate. The maximum estimate among multiple predictions for such cases is taken as the final prediction, during both training, (i.e. when computing the log-loss), and testing. The architecture is illustrated in Figure 2b.

#### SMILES convolutional network (SMILESConv)

Convolutional neural networks (CNN) are known to provide a powerful way of automatically learning complex features in vision tasks, see e.g. [12]. More recently, convolutional networks have also been shown to be effective for modeling sequential data, such as natural texts, see e.g. [30]. SMILESConv architecture is built upon 1D convolutional operators for representation learning on the SMILES strings. In this case, the kernels are vectors and they learn to leverage the relations across the consecutive characters.

Our network contains 200 1D-convolutional layers where the kernel sizes range from 1 to 200. Each layer has 32 output channels, which are followed by batch normalization [10]. We use ReLU activation function and max-pooling operators. The size of the pooling operations is equal to size the feature map that has been extracted after convolution, batch normalization, and ReLU operations. Each vector is concatenated to pass through classification layers. The extracted feature vector has 6400 units (32×200). We use dropout with a drop probability of 0.2 before the fully connected classification layers. The classification block contains 2000 units. Batch normalization and ReLU activation follow each fully connected layer. The sigmoid activation function is applied to the output layer. The overall SMILESConv architecture is shown in Figure 3.

**Fig. 3:**
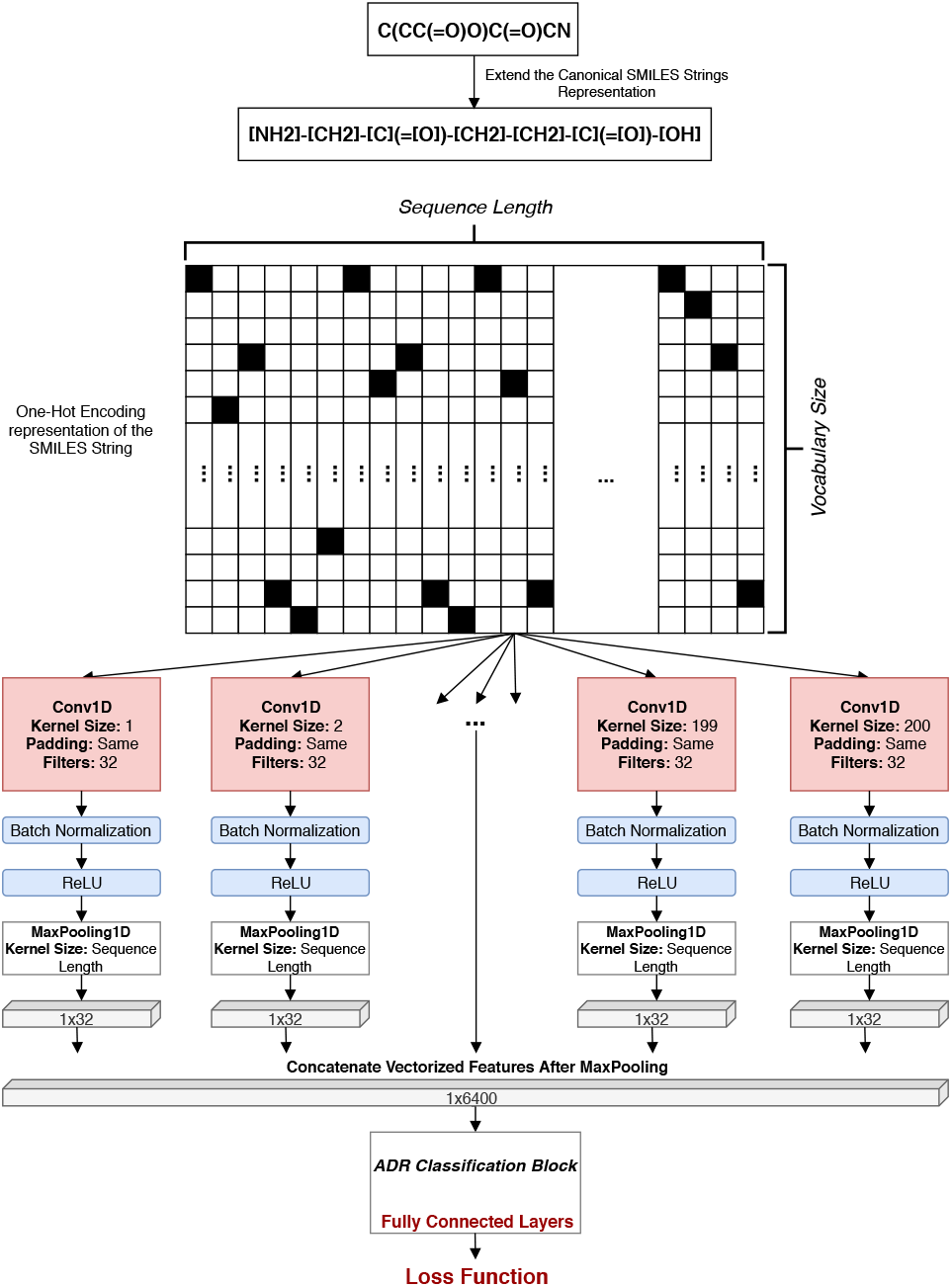
The SMILESConv architecture that performs 1D convolution operations on the SMILES representations of drugs. Fused embeddings are fed into a fully connected multi-layer classification block.

## 3 Results

### 3.1 Experimental Setup

We use 3-fold cross-validation to evaluate our models; the folds are stratified based on drugs. That is, all experiments of a single drug are either completely in the training set or completely in the test set, and therefore, a model is expected to predict the side-effects of previously unseen drugs at test time. To accomplish a fair comparison among models, we use 6 different data settings. The first 3 settings consider 791 drugs and are used to train and test only the MLP based models. The first setting uses all ~18k experiments conducted for the 791 drugs in different cell lines, dosages and time points. In this setting, each instance is an experiment for a drug and can accompany chemical structure information. The training data contains ~12k instances, while the test data contains ~6k instances. The second setting covers only the *highest quality* experiment for each of the 791 drugs, as marked in the meta-data of the LINCS L1000 dataset. Again, each instance is an experiment for a drug. The training data contains 528 instances and the test data contains 263 instances. Note that this setting is same as the one used in [32]. The third setting uses a mixture of the first two ones: ~12k instances are used for training, and 263 highest quality experiments are used for testing.

The last three settings use the 615 drugs (out of 791) which are selected according to the SMILES string criteria described in Section 2.2. To make a fair comparison between the SMILESConv and MLP based models, we re-evaluate the MLP based models in these settings and choose the best performing one to compare against SMILESConv. The fourth setting uses only the CS or SMILES string features and uses 410 samples for training and 215 samples for testing. The fifth setting uses ~9K experiments from the GEX dataset for the 410 drugs used for training and ~4K experiments for the 205 drugs for testing. Again, each instance is an experiment for a drug and can accompany CS information. The sixth setting also uses ~9K experiments from the GEX dataset for the 410 drugs like but the test data includes only the highest quality experiments of the 205 drugs.

We use binary cross entropy (BCE) as the loss function. We investigate the benefit of employing weighted BCE (WBCE) on the SMILESConv model to address the imbalance in our dataset (i.e., some side effects are observed rarely.) Adam optimizer is used for training the neural networks. While the initial learning rate for Adam optimizer is tuned separately for each model and dataset pair, the same set of hyper-parameters is used across the folds.

To assess how well we predict the side-effects of drugs overall, we use the micro-averaged Area Under Curve (AUC), micro-averaged mean Average Precision (mAP) and Hamming loss metrics. To evaluate per side effect prediction performance, we use the macro-averaged Area Under Curve (AUC) and macro-averaged mean Average Precision (mAP) metrics.

### 3.2 Performances of DeepSide Architectures

We present MLP-based model results in Table 1. Our first finding is that the base MLP model that uses only the CS fingerprints outperforms the state of the art model [32], which uses the same CS fingerprints along with the GO-transformed GEX dataset, in terms of both micro and macro AUC scores. Note that this comparison is based on the same set of drugs and side effects.

**Table 1:** Performance comparison between MLP models that use GEX, CS and META information. *X*&*Y* represents the independent two datasets that are used as inputs for the MMNN architecture. *X* is an input for one of the branches and *Y* is the input for the other branch of the MMNN-based models. [*X, Y*] represents the concatenated features of the *X* and *Y* datasets. *FC neurons* column denotes neuron size in the fully connected layers. *layers* column states the number of fully connected layers in the feature extractor and classification parts of the network.

| Model | Features | # train | # test | FC neurons | layers | Macro AUC | Micro AUC | Macro mAP | Micro mAP | Hamming |
| --- | --- | --- | --- | --- | --- | --- | --- | --- | --- | --- |
| Wang et al. [32] | GO+CS | 528 | 263 | - | - | 0.679 | 0.854 | - | - | 0.083 |
| MLP | CS | 528 | 263 | 800 | 3 - 1 | 0.784 | 0.866 | **0.457** | 0.578 | 0.072 |
| MLP | CS | 528 | 263 | 2000 | 3 - 1 | 0.781 | 0.860 | 0.454 | 0.557 | 0.075 |
| ResMLP | CS | 528 | 263 | 800 | 101 - 1 | 0.768 | 0.843 | 0.428 | 0.520 | 0.077 |
| MLP | GEX | 12K | 6K | 800 | 3 - 1 | 0.621 | 0.781 | 0.382 | 0.491 | 0.203 |
|  |  |  | 263 |  |  | 0.674 | 0.801 | 0.401 | 0.498 | 0.176 |
| MLP | [GEX, CS] | 12K | 6K | 2000 | 5 - 1 | 0.761 | 0.838 | 0.404 | 0.538 | 0.089 |
|  |  |  | 263 |  |  | 0.768 | 0.841 | 0.411 | 0.541 | 0.081 |
| MLP | [GEX, CS, META] | 12K | 6K | 2000 | 5 - 1 | 0.767 | 0.845 | 0.421 | 0.558 | 0.086 |
|  |  |  | 263 |  |  | 0.774 | 0.844 | 0.426 | 0.528 | 0.076 |
| MLP | [GEX, CS, META] | 528 | 263 | 2000 | 5 - 1 | 0.727 | 0.832 | 0.390 | 0.497 | 0.089 |
| ResMLP | [GEX, CS, META] | 12K | 6K | 2000 | 5 - 1 | 0.760 | 0.857 | 0.422 | 0.577 | 0.084 |
|  |  |  | 263 |  |  | 0.771 | 0.856 | 0.428 | 0.547 | 0.075 |
| MTNN | [GEX, CS, META] | 12K | 6K | 2000 | 5 - 1 | 0.759 | 0.841 | 0.401 | 0.511 | 0.087 |
|  |  |  | 263 |  |  | 0.772 | 0.851 | 0.418 | 0.522 | 0.079 |
| MMNN.Sum | CS & [GEX, META] | 12K | 6K | 800 - 800 | 3 - 1 | 0.772 | 0.871 | 0.435 | 0.600 | 0.081 |
|  |  |  | 263 |  |  | **0.790** | **0.877** | **0.457** | 0.592 | **0.070** |
| MMNN.Concat | CS & [GEX, META] | 12K | 6K | 800 - 800 | 3 - 1 | 0.768 | 0.868 | 0.436 | 0.598 | 0.080 |
|  |  |  | 263 |  |  | 0.787 | 0.875 | 0.460 | 0.586 | 0.072 |
| MMNN.Sum | CS & GEX | 12K | 6K | 800 - 800 | 3 - 1 | 0.764 | 0.868 | 0.431 | **0.602** | 0.081 |
|  |  |  | 263 |  |  | 0.779 | 0.872 | 0.445 | 0.582 | 0.071 |
| MMNN.Sum | CS & GEX | 12K | 6K | 2000 - 2000 | 3 - 1 | 0.772 | 0.864 | 0.440 | 0.588 | 0.082 |
|  |  |  | 263 |  |  | 0.783 | 0.867 | 0.444 | 0.569 | 0.073 |
| MMNN.Sum | CS & GEX | 528 | 263 | 2000 - 2000 | 3 - 1 | 0.772 | 0.863 | 0.424 | 0.557 | 0.075 |

The MLP model based purely on GEX features yields the lowest scores in both Settings 1 and 3 (Table 1), the macro-AUC is at most 67% and macro-AUC is 80%. This indicates that GEX features alone are not sufficiently informative for side effect prediction. When we combine GEX and CS features through concatenation and input to the MLP model, the performance increases to 76.8% macro AUC and 84.1% micro AUC scores (Setting 3; similar for Setting 1), which are still below the performance of the MLP model trained only with the CS features.

The ResMLP architecture, which uses residual connections across the fully connected layers does not improve upon the base MLP model. MTNN, which aims to leverage the side effect group information based on the side effect ontology, does not improve over the base MLP model either. On the other hand, the MMNN model, which uses two modalities (one for the concatenated GEX profiles and META information and the other for the CS fingerprints), produces the best predictive performance among all MLP-based architectures in terms of all metrics, with the exception of micro mean average precision (micro mAP). This architecture achieves 0.111 macro AUC improvement and 0.023 micro AUC improvement over state of the art in Setting 3 when summation based embedding fusion is used. Concatenation based fusion yields similar results. MMNN is the only architecture that benefits from adding GEX features on to the CS features. Since we consistently obtain very similar or better results by incorporating the META information, we exclude the results of some of the models without META features for brevity.

Setting 2 only uses the highest quality experiments (as in [32]), whereas Setting 3 uses the all experiments for a compound during training. For testing, both settings use the highest quality experiments. Here, we validate our hypothesis that a deep learning framework should be able to perform better by utilizing the full dataset in Setting 3. First, we compare the performance of the MLP model under Setting 2 and Setting 3 (using GEX, CS, and META features): using Setting 3 provides 4.7% macro AUC and 1.2% micro AUC, 3.1% macro mAP and 3.6% micro mAP improvement over Setting 2. We also compare the performance of the best MLP-based model (MMNN.Sum) under these two settings using the CS and GEX features. Indeed, setting 3 provides 1.1% macro AUC, 0.4% micro AUC, 2.4% macro mAP and 1.2% micro mAP improvement over Setting 3. While the margin of improvement is smaller for the more complex model, both results show the benefit of using all experiments in the LINCS L1000 dataset.

We investigate the benefit of using SMILES strings for representing drug structures and employing convolutional neural networks to extract features on them. Table 2 shows the results of SMILESConv models that are trained with unweighted (BCE) and class weighted loss (WBCE) functions. To make a fair comparison to the SmilesConv models, we retrain separate MLP and MMNN.Sum architectures with datasets of Settings 4 - 6. In SMILESConv models, cost-sensitive training with WBCE improves the results compared to training with BCE; all performance measures are higher for WBCE except for the hamming loss. SMILESConv outperforms both the MMNN.Sum and the MLP based model; with WBCE, it achieves 0.809 macro AUC and 0.885 micro AUC. This corresponds to about 2.1% improvement in macro AUC and 3.6% improvement in micro AUC compared to the MLP model that uses only the CS structures. It also improves upon the MMNN.Sum about 1.5% in macro AUC and 3.3% in micro AUC. Similar improvements are observed for the other performance metrics MAP and Hamming loss. The predicted probabilities by SMILESConv WBCE for every compound - side effect pair are listed in Supplementary Table 1.

**Table 2:**
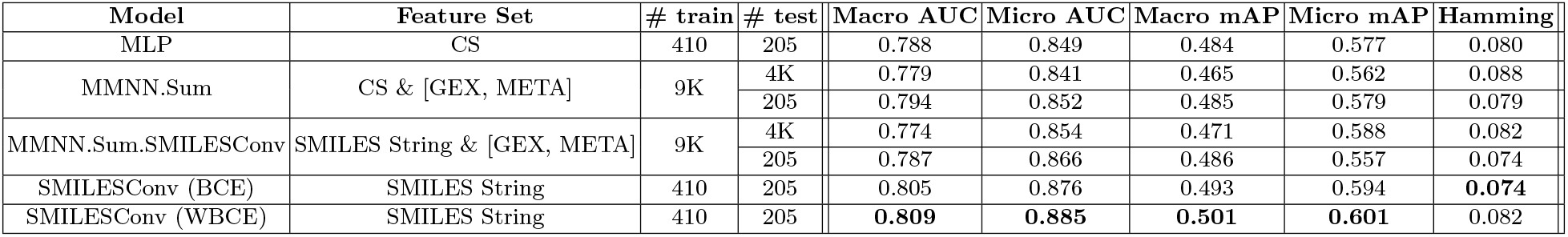
Performance comparison between MLP and Conv models which are trained with 615 drugs for the 1042 side effects. [*X, Y*] represents the concatenated features of the *X* and *Y* datasets. [*X*]&[*Y*] represents the two separate datasets applied different braches of the MMNN-based models. BCE denotes binary cross entropy and WBCE denotes the weighted binary cross entropy.

| Model | Feature Set | # train | # test | Macro AUC | Micro AUC | Macro mAP | Micro mAP | Hamming |
| --- | --- | --- | --- | --- | --- | --- | --- | --- |
| MLP | CS | 410 | 205 | 0.788 | 0.849 | 0.484 | 0.577 | 0.080 |
| MMNN.Sum | CS & [GEX, META] | 9K | 4K | 0.779 | 0.841 | 0.465 | 0.562 | 0.088 |
|  |  |  | 205 | 0.794 | 0.852 | 0.485 | 0.579 | 0.079 |
| MMNN.Sum.SMILESConv | SMILES String & [GEX, META] | 9K | 4K | 0.774 | 0.854 | 0.471 | 0.588 | 0.082 |
|  |  |  | 205 | 0.787 | 0.866 | 0.486 | 0.557 | 0.074 |
| SMILESConv (BCE) | SMILES String | 410 | 205 | 0.805 | 0.876 | 0.493 | 0.594 | **0.074** |
| SMILESConv (WBCE) | SMILES String | 410 | 205 | **0.809** | **0.885** | **0.501** | **0.601** | 0.082 |

We further investigate whether uniting the GEX and META features with SMILES strings improves the performance. We train a new MMNN.Sum model in which we replace the chemical fingerprint representation (CS) with the SMILES representation. We observe that this improves MMNN.Sum model’s performance but cannot outperform SMILESConv only model (Table 2).

## 4 Discussion

We investigate the easiest (top-10) and the hardest (bottom-10) side effects to predict by the SMILESConv model (WBCE) in Table 3. For both cases, these side effects have less than 100 positive samples. Although there is no clear pattern, we observe that the easy examples are relatively more specific compared to the hard examples (i.e., Myocardial rupture, Lupus miliaris disseminatus faciei, and Paraplegia vs. Ear disorder, Personality Disorder and Sensory disturbance). We also investigate our most confident but incorrect predictions. Table 3 shows the top-10 false positive and top-10 false negative predictions. For the following false positive examples, we find evidence in the literature that the predicted side effects might be relevant. Daunorubicin, which is a chemotherapeutic compound, is predicted to cause anemia by DeepSide. Chemotherapy-induced anemia is a common side effect in cancer patients [6]. In particular for this drug, Hazardous Substances Data Bank^5^ (a toxicology database curated by NIH NLM Toxicology Network) lists anemia as a possible adverse reaction for daunorubicin^6^. Similarly, we find that sulfasalazine (a drug used to treat rheumatoid arthritis and ulcerative) causes vomitting. This finding is supported by [21], which reports that 64 out of 152 people developed adverse reactions due to this drug, and 19 out of that 64 had vomiting. Finally, our model predicts halcinonide to cause hypertension. Halcinonide is a corticosteroid that is used to treat various skin conditions. It is a glucocorticoid and [18] lists hypertension as an adverse effect for glucocorticoids. Note that none of the above findings are reported in SIDER. We also find support for 9 out of the top 10 false positives through commercial online resources. Nevertheless, it is hard to assess the reliability as there is no peer review system. While it is harder to evaluate false negatives, we find that rather than (i) doxycycline causing premenstrual syndrome, and (ii) cyproterone causing leiomyoma; they are used in the treatment of these conditions [29, 24]. For the rest of the findings we see that there are indications in the literature and commercial online resources that these compounds cause corresponding side effects.

**Table 3:** The easiest (top-10) and the hardest (bottom-10) side effects to predict by the SMILESConv model trained with the weighted binary cross-entropy loss. Number of positive samples column indicates the number of drugs annotated with a given side effect.

| Side Effect | # Pos. Samples | SMILESConv AUC |
| --- | --- | --- |
| Skin test positive | 21 | 1.00 |
| Cushing’s syndrome | 12 | 1.00 |
| Myocardial rupture | 19 | 1.00 |
| Alkalosis hypokalaemic | 21 | 1.00 |
| Fat embolism | 15 | 1.00 |
| Muscle mass | 21 | 1.00 |
| Coombs direct test positive | 10 | 1.00 |
| Paraplegia | 17 | 1.00 |
| Lupus miliaris disseminatus faciei | 23 | 1.00 |
| Nitrogen balance | 23 | 1.00 |
| Skin burning sensation | 7 | 0.54 |
| Panic attack | 9 | 0.55 |
| Tachypnoea | 10 | 0.64 |
| Sensory disturbance | 8 | 0.50 |
| Hepatitis fulminant | 11 | 0.58 |
| Ear disorder | 28 | 0.57 |
| Arrhythmia supraventricular | 15 | 0.65 |
| Respiratory disorder | 87 | 0.64 |
| Personality disorder | 26 | 0.62 |
| Congenital eye disorder | 11 | 0.62 |

**Table 4:**
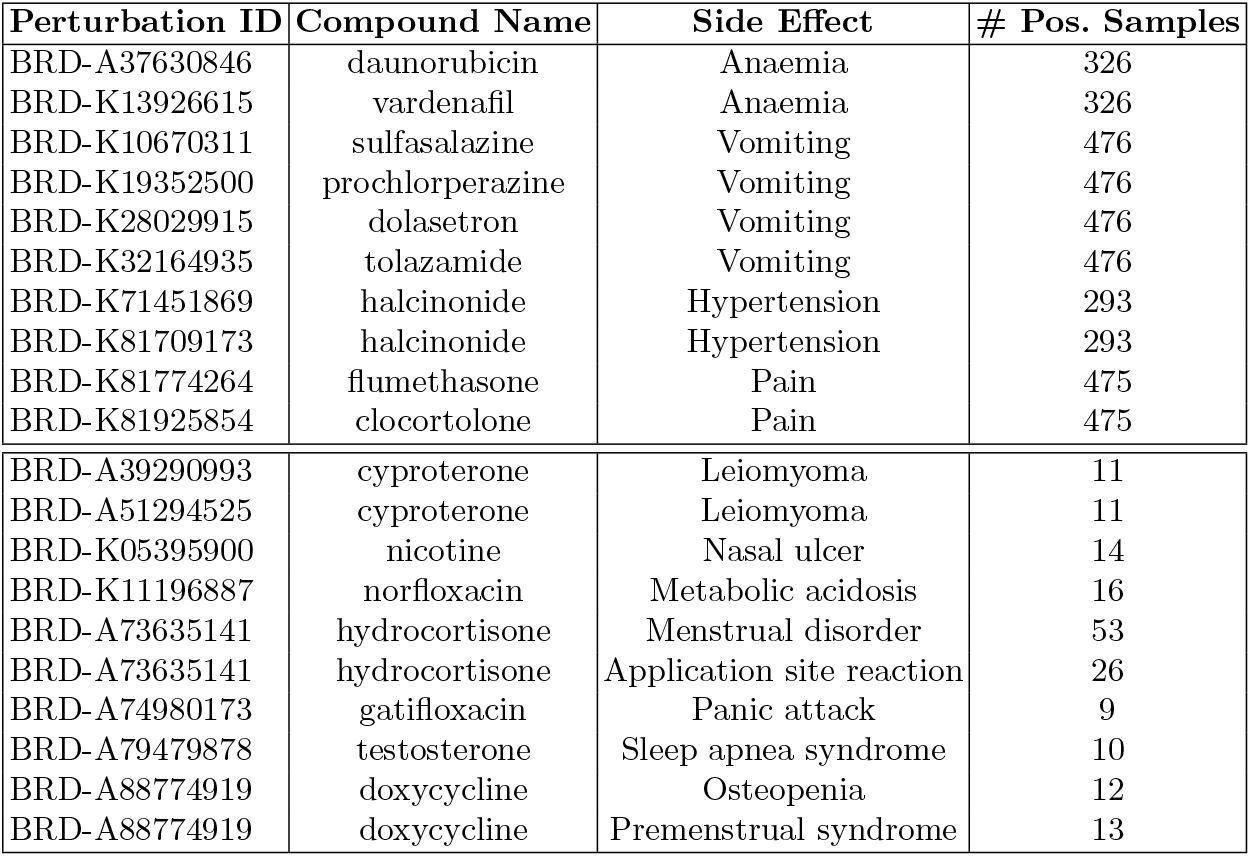
The top-10 false positive (top table) and top-10 false negative predictions (bottom table) of the SMILESConv WBCE model. These are the most confident predictions by the model that are contradicting with the ground truth. For the listed false positive pairs, predicted probabilities are > 0.9995. For the false negative pairs, predicted probability scores are < 0.0005. Note that a drug (name) might have multiple pertubagen id that correspond to different SMILES strings. In that case drug name - side effect pairs are listed multiple times.

The LINCS L1000 dataset is a useful resource for predicting condition specific side effects. In our experiments though, we find the GEX does not improve the results substantially (see Tables 1 2) and the best performing model that relies on the drug structure and surpasses the state-of-the-art performance [32] (see Tables 1 and 2). One reason for not being able to leverage condition specific GEX information despite employing various deep learning architectures could be the absence of the condition specific ground truth labels. Since the available side effect labels are per drug but not per condition-drug pairs (i.e., dosage - drug), we suspect the model cannot make use of the LINCS dataset as effectively as it could. On the other hand, deep learning framework can leverage the chemical structure information well and can surpass state of the art result, which uses chemical structure and gene expression features in combination with gene ontology [32].

## 5 Conclusion

The pharmaceutical drug development process is a long and demanding process. Unforeseen ADRs that arise at the drug development process can suspend or restart the whole development pipeline. Therefore, the a priori prediction of the side effects of the drug at the design phase is critical.

In our DeepSide framework, we use context-related (gene expression) features along with the chemical structure to predict ADRs to account for conditions such as dosing, time interval, and cell line. The proposed MMNN model uses GEX and CS as combined features and achieves better accuracy performance compared to the models that only use the chemical structure (CS) fingerprints. The reported accuracy is noteworthy considering that we are only trying to estimate the condition-independent side effects. Finally, SMILESConv model outperforms all other approaches by applying convolution on SMILES representation of drug chemical structure.

5 http://toxnet.nlm.nih.gov/cgi-bin/sis/htmlgen?HSDB

6 http://toxnet.nlm.nih.gov/cgi-bin/sis/search2/r?dbs+hsdb:@term+@rn+@rel+20830-81-3, accessed Oct 30, 2019

